# Individual-Level Associations of Cognition and Personality With Adolescent Mental Health Vary Across Brain and Environmental Profiles

**DOI:** 10.64898/2025.12.25.696521

**Authors:** Jiadong Yan, Bin Wan, Paule Joanne Toussaint, Judy Chen, Gleb Bezgin, Yasser Iturria-Medina, Danilo Bzdok, Alan C. Evans, Sherif Karama

**Affiliations:** Montreal Neurological Institute, McGill University, Montreal, Canada; McGill Centre for Integrative Neuroscience, McGill University, Montreal, Canada; Department of Psychiatry, University Hospitals of Genève, Thonex, Switzerland; Max Planck Institute for Human Cognitive and Brain Sciences, Leipzig, Germany; Institute of Neuroscience and Medicine, Research Center Jülich, Jülich, Germany; Ludmer Centre for NeuroInformatics and Mental Health, Montreal, Canada; Mila - Quebec Artificial Intelligence Institute, Montreal, Canada; Douglas Mental Health University Institute, McGill University, Montreal, Canada

## Abstract

Associations between psychopathology, cognition, and personality are well established. However, these associations unfold within a broader developmental context shaped by brain maturation, environmental exposures and pubertal transitions, whose influences remain poorly understood. Here, we characterized how cognition and personality relate to adolescent mental health within this developmental context. Using data from the Adolescent Brain Cognitive Development Study (*N* = 4501), we simultaneously modeled 16610 brain structural, environmental, and pubertal measures to examine 165 behavioral pairs comprising cognitive and personality traits with mental health symptoms. Using an integrated analytic pipeline combining multifactorial mediation and longitudinal interaction-inclusive models, we evaluated whether developmental changes in cognition and personality relate to mental health at the individual level, and whether these associations vary across brain, environmental, and pubertal profiles. Most behavioral pairs showing significant mediation effects likewise exhibited significant longitudinal effects. These pairs were primarily influenced by neurobiological and environmental measures; cortical frontal and temporal regions relevant for cognitive control and emotional regulation as well as family and lifestyle contexts were most prominently implicated. Finally, after accounting for inter-individual heterogeneity, personality traits emerged as the main behavior associated with mental health. These findings characterize developmental associations linking cognition and personality with adolescent mental health, and show that these associations vary across brain, environmental, and pubertal profiles. The study advances understanding of behavioral development and provides a framework for identifying targets for psychopathology prevention.

## INTRODUCTION

Adolescence is a developmental period during which many psychiatric disorders first emerge (1,2). During this time, cognitive abilities and personality traits also undergo substantial developmental changes (3–5). Numerous studies have reported stable associations between cognition, personality, and mental health symptoms in youth (6–9). For example, lower cognitive performance has been linked to increased risk of mood and anxiety disorders (10). Likewise, all Big Five personality traits have been associated with well-being (11). However, despite these well-documented associations, the developmental mechanisms linking cognition and personality to mental health remain poorly understood (12,13).

Well-established developmental theoretical frameworks (14–16), emphasize that behavioral development reflects dynamic interactions among brain maturation, environmental exposures, and pubertal transitions. Previous studies have examined relationships among biological, environmental and behavioral factors, but have typically focused on a limited subset of these domains (17–19). Consequently, studies linking cognition and personality to mental health have rarely considered these developmental influences jointly (20–22). This limitation may lead to incomplete characterization of behavioral relationships and hinder mechanistic understanding of how cognitive and personality traits relate to psychopathology.

Furthermore, adolescence is characterized by substantial inter-individual heterogeneity in mental health and behavioral development (23,24). Differences in neurobiology (19,25), environmental exposures (26,27), and pubertal maturation (28,29) contribute to this variability. These findings suggest that associations of cognitive and personality traits with mental health are unlikely to be uniform across individuals. Although recent psychopathology research has attempted to move beyond traditional group-level analyses using subtyping approaches, resulting inferences remain at the subgroups rather than the individual level (23,30). As a result, how behavioral associations vary across individuals and how such variability relates to differences in brain, environmental, and pubertal profiles, remains largely unknown.

To address these gaps, we investigated individualized associations of cognition and personality with adolescent mental health within an integrated developmental framework incorporating brain, environmental, and pubertal factors (**Figure 1B-D**). Leveraging data from 4501 participants in the Adolescent Brain Cognitive Development (ABCD) Study across baseline and follow-up assessments, we analyzed 165 behavioral associations connecting 6 cognitive traits and 9 personality measures with 11 mental health symptoms while jointly considering 16610 biological and environmental measures (17,31). Specifically, we first identified pathways linking biological and environmental factors with cognition, personality, and mental health. We then examined whether longitudinal changes in cognition or personality were associated with corresponding changes in mental health symptoms and whether these relationships varied across brain, environmental, and pubertal profiles. Finally, we characterized how these behavioral associations vary across individuals.

**Figure 1.**
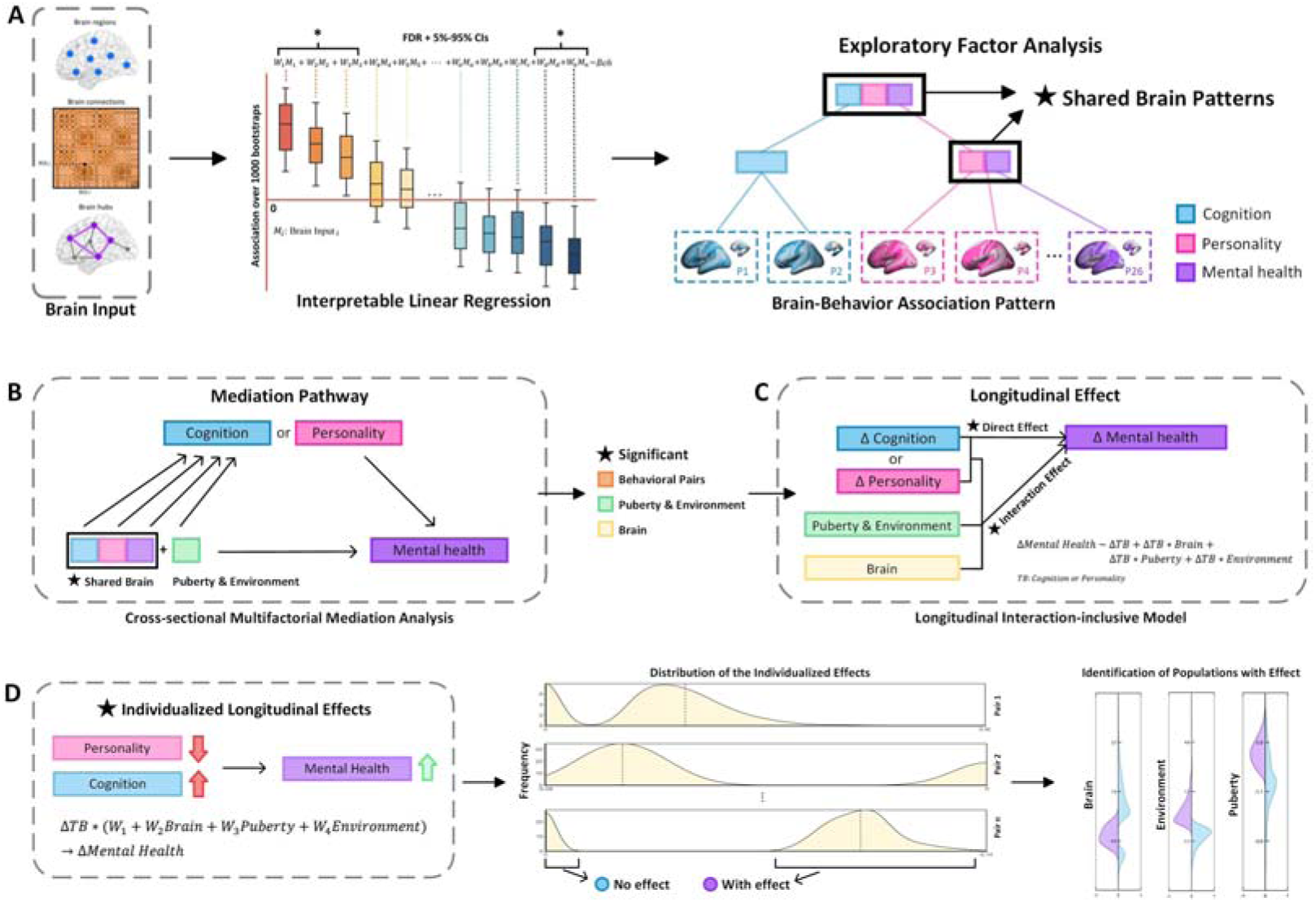
Flowchart of the analysis. **(A)** Brain structural features were first associated with cognition, personality, and mental health measures to identify brain-behavior association patterns. Exploratory factor analysis was then applied to identify shared brain patterns between cognition/personality and mental health. **(B)** Based on the shared brain features identified, cross-sectional multifactorial mediation analysis was used to identify pathways linking cognition or personality to mental health through brain, environmental, and pubertal factors. **(C)** Based on the factors involved in the identified pathways, longitudinal interaction-inclusive models examined whether changes in cognition or personality were associated with corresponding changes in mental health and whether these links were shaped by brain, environmental, and pubertal profiles. **(D)** Individualized effects derived from the longitudinal models were used to characterize heterogeneity across individuals and to identify populations showing significant behavioral effects.

## METHODS AND MATERIALS

### Participants

This study used data from the ABCD Study 5.1 data release, including phenotypic assessments collected at baseline and follow-up visits across 22 sites (32). The ABCD Study was approved by the central Institutional Review Board at the University of California, San Diego, and written informed consent was obtained from all participants. After quality control procedures (33), the final sample included 4501 participants (2207 females) for cross-sectional analyses and 1437 participants (740 females) for longitudinal analyses (see **Table S1** and the **Supplement** for details).

### Behavioral Measures

Behavioral measures comprised 6 cognitive, 9 personality and 11 mental health assessments widely used in prior research (7,31,34,35). Cognitive abilities included 6 scores from the NIH Toolbox (36) and the Little Man Test (37), the only cognitive measures available across all timepoints. Personality traits were assessed using 9 impulsivity-related measures from the Modified UPPS-P for Children (38) and the Behavioral Inhibition and Activation Scales (39). Mental health measures included 11 scales from the Achenbach Child Behavior Check List (40), the Parent General Behavior Inventory (41), and the Pediatric Psychosis Questionnaire (42). More details are described in the **Supplement**.

### Brain Structural, Environmental, and Pubertal Measures

We adopted a comprehensive set of brain structural measures, as in a previous study (31). Structural and diffusion MRI data were obtained from the ABCD Study and preprocessed by the ABCD consortium (32,33). Brain parcellation was performed using the Destrieux atlas (43) for the cortex (148 regions of interest [ROIs]) and the ASEG atlas (44) for subcortex (14 ROIs). For each brain ROI, we extracted 6 structural MRI, 4 diffusion tensor imaging, and 6 restriction spectrum imaging measures (45). Morphometric similarity networks (46) were then constructed to characterize structural connections between ROIs, and 6 graph-theoretical metrics (47) were computed for each ROI to derive hub-wise measures. In total, 16563 brain structural features were included in the analyses (see the **Supplement** for details). Moreover, we included 39 environmental measures across 5 categories (17): 13 perinatal and early development events, 7 life events and lifestyle factors, 10 family variables (48), 6 neighborhood variables, and 3 school variables. For puberty, we adopted 8 measures (14) consisting of 3 domains: 4 Pubertal Development Scale measures (49,50) capturing puberty stage, Body Mass Index, and 3 hormone measures (49,51). The detailed measures are described in the **Supplement**.

### Identification of Informative Brain Features

To reduce the dimensionality of brain features and mitigate issues related to the sample-to-predictor (N/P) ratio (52) in subsequent machine learning analyses, we identified the most informative brain structural features (**Figure 1A**). Briefly, for each of the 26 behavioral measures, we first fitted separate interpretable linear regression models (31) to estimate brain-behavior association patterns across 16563 brain structural features. We then applied exploratory factor analysis (53) to these association maps to identify shared brain patterns across behaviors (**Figure S1**). Based on these shared patterns, we extracted the most informative subset of brain features according to the magnitude of their loadings on the shared patterns, which were used in the subsequent analyses. A detailed description of the analysis is provided in the **Supplement**.

### Cross-Sectional Multifactorial Mediation Analysis

To examine whether cognition or personality statistically mediated the associations between brain structural, environmental, and pubertal factors and mental health outcomes, we developed a multifactorial mediation framework applied to cross-sectional data (**Figure 1B**). For each of the 165 behavioral pairs, one cognition or personality measure was specified as the mediator and one mental health measure as the outcome. Reduced brain structural measures together with environmental and pubertal variables were jointly entered as predictors (see **Figure S2** and the **Supplement** for details). This framework extends conventional mediation models by allowing multiple predictors to be evaluated simultaneously rather than testing a single predictor (54). Model parameters were estimated using linear regression with 1000 bootstrap resampling iterations, and effects were considered significant when the two-sided 5-95% bootstrap confidence intervals excluded zero (55–58). Additional methodological details are provided in the **Supplement**. Behavioral pairs showing significant mediation effects, together with the corresponding significant brain, environmental, and pubertal variables, were retained for subsequent longitudinal analyses.

### Longitudinal Interaction-Inclusive Model

To examine whether longitudinal changes in cognition or personality were associated with corresponding changes in mental health, and whether these associations were modulated by brain, environmental, and pubertal profiles, we developed a longitudinal interaction-inclusive model (**Figure 1C**). For each behavioral pair, the change in the cognition or personality measure was modeled as a predictor of the change in the corresponding mental health outcome. To assess whether this longitudinal association was modulated by brain structural, environmental, and pubertal measures, interaction terms between the cognition or personality change and the baseline levels of these measures were also included as predictors. Additional methodological details are provided in the **Supplement**. Model estimation and significance testing followed the same bootstrap procedure described in the mediation analyses. Behavioral pairs showing significant longitudinal effects, together with the corresponding significant brain structural, environmental, and pubertal modulators, were retained for subsequent individualized analyses.

### Individualized Analysis

To characterize inter-individual variability in the identified behavioral links, we performed individualized analyses based on the interaction terms derived from the previous longitudinal models (**Figure 1D**). For each participant, we estimated an individualized effect representing the predicted change in mental health associated with a one-standard deviation change in the corresponding cognition or personality measure, conditional on the individual’s brain, environmental, and pubertal profile. Using 1000 bootstrap resamples, two-sided 5-95% confidence intervals were estimated for each individualized effect. Individuals whose confidence intervals excluded zero were considered to show significant effects. We then quantified the proportion of such individuals across the sample and estimated the mean effect among them. Finally, baseline brain, environmental, and pubertal measures were compared between individuals with and without significant effects to identify profiles associated with a higher likelihood of exhibiting significant behavioral effects. Additional methodological details are provided in the **Supplement**.

### Statistical Controls and Multiple-Comparison Correction

To reduce collinearity among predictors and ensure interpretable regression estimates, we orthogonalized predictors prior to model fitting using a principal component transformation.

Age, sex, handedness, ethnicity, study site, and brain volume were included as covariates in all analyses. For the longitudinal analyses, we additionally controlled for changes in the time-varying variables included in the models. Subject and family confounds were further addressed during the bootstrap procedure (31).

Because multiple behavioral pairs were tested, multiple-comparison correction was applied across models using both empirical and competitive permutations (59). Correction within each model was unnecessary because all input coefficients were jointly estimated within a single linear regression framework (55). Additional methodological details are provided in the **Supplement**.

### Model Validation

To assess model generalizability, split-half validation was performed. For each behavioral pair, the sample was randomly divided into two independent subsets of equal size, and the same modeling procedure was applied to both subsets. Significant parameter estimates from the two models were compared using the Pearson correlation coefficient. This process was repeated 100 times, and the mean correlation across repetitions was used as an indicator of model generalizability.

### Code and Data Availability

Data used in this study were obtained from the ABCD Study (release 5.1). All analysis code is accessible online (https://github.com/JDYan/Behavioral-Causality).

## RESULTS

### Multifactorial Mediation Linking Cognition and Personality to Mental Health

Among the 165 behavioral pairs tested, 95 exhibited significant mediation effects, with the proportions mediated reaching up to 19.88% (**Figure 2A**). Personality showed greater involvement in these mediation pathways than cognition, particularly through behavioral inhibition and negative urgency. Crystallized cognitive abilities, including vocabulary and reading, were also frequently involved. Among mental health outcomes, attention problems, rule-breaking behavior, and psychosis symptoms and severity were most frequently implicated.

**Figure 2.**
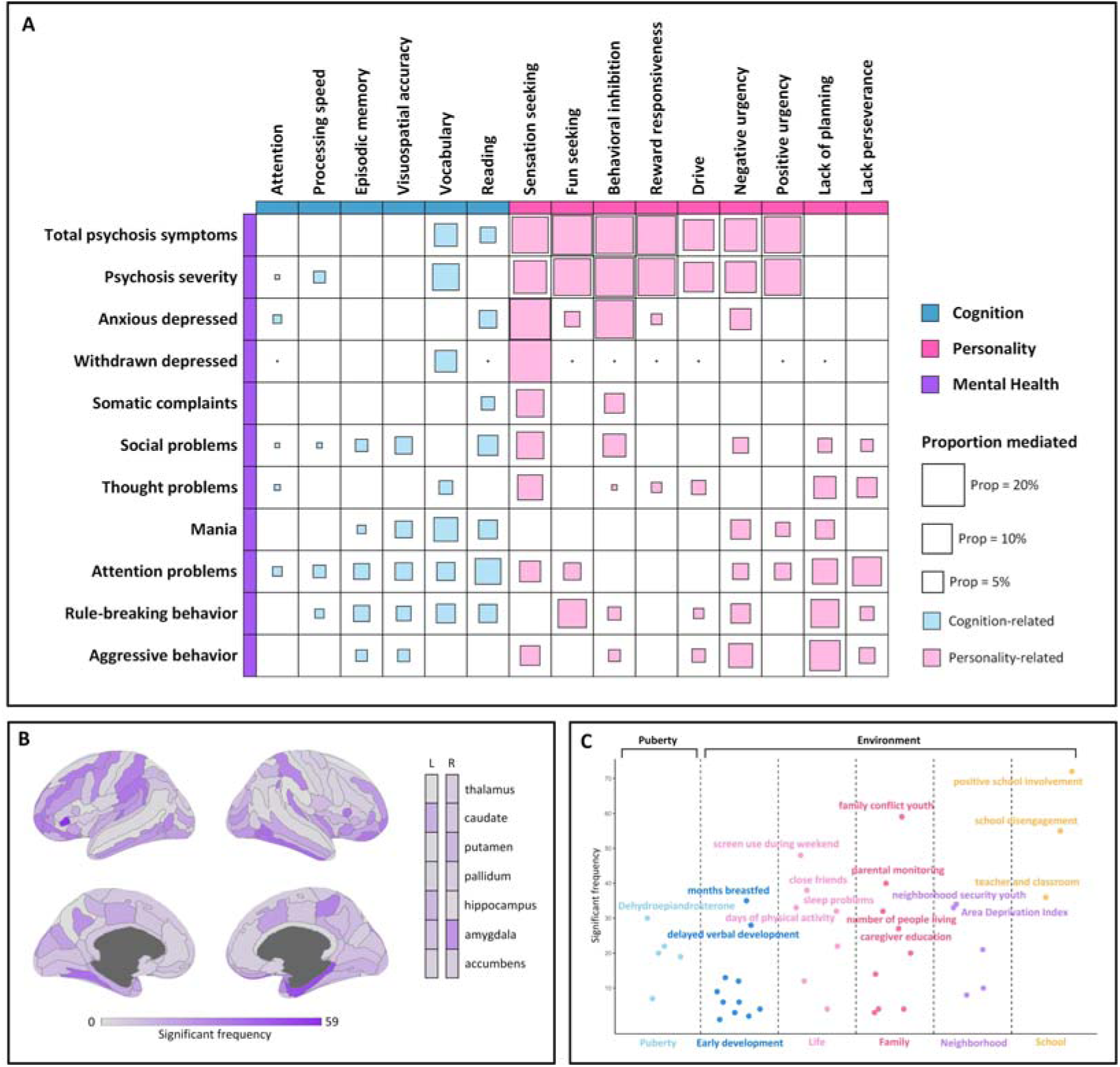
Brain, environmental, and pubertal influences on adolescent mental health mediated by cognition and personality. **(A)** Proportion mediated for each behavioral pair. Square size represents the proportion mediated, with blue indicating cognition-related pairs and pink indicating personality-related pairs. **(B)** Summary of brain regions significantly implicated in the mediation pathways. Across all 95 significant behavioral pairs, significant brain regions, connections, and hubs were counted based on 162 ROIs; each connection was counted as two ROIs to reflect that each connection contributes to the involvement of both linked regions. **(C)** Summary of pubertal and environmental factors contributing to the mediation pathways. For all 95 significant behavioral pairs, we quantified the frequency of significant pubertal and environmental measures, with the most prominent ones displayed.

Brain structure contributed to most mediation pathways. The most frequently implicated regions were located in frontal (left horizontal ramus of the anterior segment of the lateral sulcus and right medial orbital sulcus), temporal (right inferior temporal gyrus and right parahippocampal gyrus), and occipitotemporal cortices (left medial occipitotemporal sulcus and lingual sulcus; **Figure 2B**). Separate summaries for brain regions, connections, and hubs are shown in **Figure S3**.

Environmental factors were significantly involved in all 95 pathways, with weekend screen time, family conflict, and school involvement and disengagement emerging as the most frequently implicated contributors (**Figure 2C**). In contrast, pubertal measures played a more limited role, with dehydroepiandrosterone (DHEA) showing relative prominence. Results with labels for all measures are provided in **Figure S4**.

### Longitudinal Effects of Cognition and Personality on Mental Health

Among the 95 behavioral pairs identified in the mediation analysis, 68 showed significant longitudinal effects in the interaction-inclusive models (**Figure 3A**). Consistent with the mediation findings, personality contributed more frequently than cognition to these longitudinal relationships. Substantial combinatorial heterogeneity was observed among four effect types across behavioral pairs (direct effects and modulation by brain, environmental, and pubertal factors), with 31 of the 68 pairs exhibiting at least two effect types. Direct effects from cognition or personality to mental health were observed in 30 pairs.

**Figure 3.**
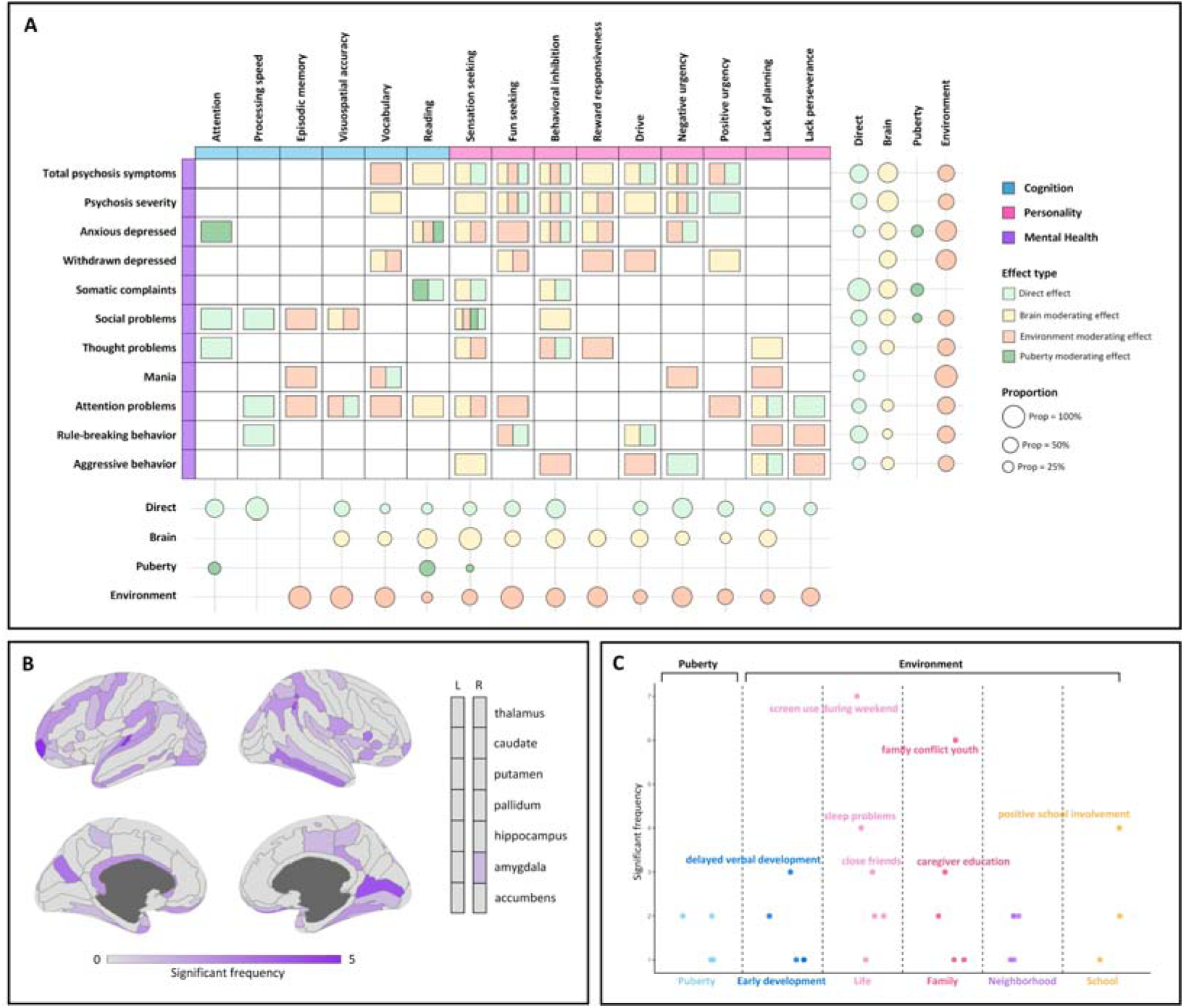
Longitudinal effects linking cognition and personality to mental health and their modulation by brain, pubertal, and environmental factors. **(A)** Types of longitudinal effects across behavioral pairs. Four effect types were examined: direct effects (light green), modulating effects by brain (yellow), puberty (green), and environment (orange). Circle size indicates the proportion of significant effects, calculated as the ratio of behavioral pairs showing a significant effect to the total number of significant pairs within each row or column. **(B)** Summary of brain regions showing significant modulation effects. Across the 68 significant behavioral pairs, significant brain regions, connections, and hubs were counted based on 162 ROIs; each connection was counted as two ROIs to reflect that each connection contributes to the involvement of both linked regions. **(C)** Summary of pubertal and environmental factors showing significant modulation effects. Across the 68 significant pairs, the frequency of significant pubertal and environmental measures was quantified, with the most prominent ones displayed.

Brain modulation was widespread, occurring in 34 of the 68 behavioral pairs. The most frequently implicated regions were located in frontal (left frontomarginal gyrus and sulcus), temporal (left transverse temporal sulcus), and occipital cortices (right sulcus intermedius primus and right calcarine sulcus; **Figure 3B**), highlighting the continued importance of frontal and temporal regions observed in the mediation results. Among subcortical structures, only the right amygdala showed consistent significance. Separate summaries for brain regions, connections, and hubs are shown in **Figure S5**.

Environmental modulation also showed substantial involvement. Compared with the mediation findings, the most prominent environmental contributors in the longitudinal models shifted from school-related variables to life-event and family factors, including weekend screen time, sleep problems, and family conflict (**Figure 3C**). Pubertal influences remained minimal. Results with labels for all measures are provided in **Figure S6**.

Split-half validation of the longitudinal models demonstrated strong reproducibility (*r* = 0.81 ± 0.13). After applying multiple-comparison correction via Bonferroni adjustment (empirical permutation, adjusted *p* < 0.05), all 68 longitudinal behavioral pairs remained significant. Results of the competitive permutation-based correction are provided in the **Supplement**.

### Individualized Associations of Cognition and Personality With Mental Health

We next examined inter-individual variability in the behavioral effects identified in the previous longitudinal models. Among the 68 behavioral pairs, 37 showed significant individualized effects in at least a subset of participants, and 23 showed significant effects in more than 50% of individuals (**Figures 4A** and **S7**). Across these 23 pairs, effect directions were consistent, with increased cognitive abilities and decreased maladaptive personality traits associated with improved mental health outcomes. Personality traits were most frequently involved, particularly behavioral inhibition and negative urgency. Among cognitive measures, attention and processing speed appeared most often. The mental health outcomes most commonly implicated were attention problems and psychosis-related symptoms and severity.

**Figure 4.**
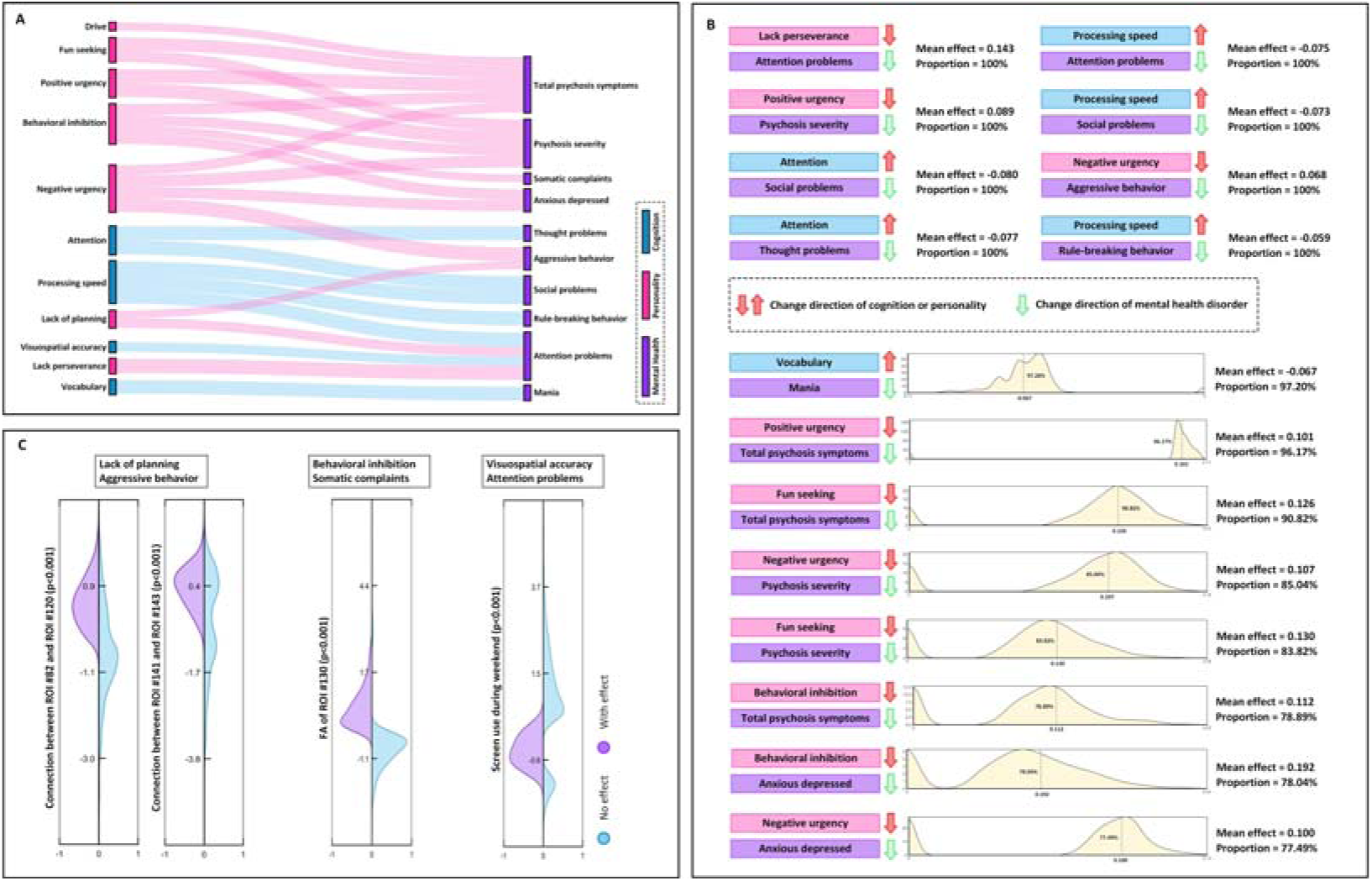
Inter-individual variability in behavioral effects linking cognition and personality with mental health. **(A)** Twenty-three behavioral pairs in which more than 50% of participants showed significant individualized effects. Blue indicates cognition-related pairs and pink indicates personality-related pairs. Ribbon width indicates the proportion of participants with significant effects. **(B)** Eight behavioral pairs showed significant individualized effects in all participants (100%). For each pair, the mean effect is shown. Red arrows indicate the direction of change in cognition or personality, whereas green arrows indicate the corresponding direction of change in the mental health outcome. Upward arrows denote increases and downward arrows denote decreases. An additional 8 behavioral pairs showed significant individualized effects in 75-100% of participants. For these pairs, the distribution of individualized effects across participants is shown together with the mean effect and the proportion with significant effects. **(C)** Examples of profiles associated with significant individualized effects. For three behavioral pairs, participants were divided into groups with and without significant individualized effects. Distributions of corresponding phenotypic profiles are shown for both groups; all differences were significant (*p* < 0.001). ROI #82: right middle-posterior part of the cingulate gyrus and sulcus; ROI #120: right marginal branch of the cingulate sulcus; ROI #130: right intraparietal sulcus and transverse parietal sulci; ROI #141: right postcentral sulcus; ROI #143: right superior part of the precentral sulcus.

We observed 8 behavioral pairs showing significant individualized effects in all participants (100%; **Figure 4B**). In contrast with the overall pattern observed in earlier analyses, cognitive traits, particularly processing speed and attention, were more frequently involved in these pairs than personality traits. Nevertheless, among the 8 pairs showing significant effects in 75-100% of individuals (**Figure 4B**), personality played a more prominent role. Across these 16 pairs, mean individualized effect sizes reached up to 0.192, and half of these pairs showed mean effects greater than 0.1. An additional 7 pairs showing significant effects in 50-75% of individuals are shown in **Figure S8**. Split-half validation of the corresponding longitudinal models for the 23 pairs (**Figure 4A**) demonstrated strong reproducibility (*r* = 0.84 ± 0.10).

To further characterize profiles associated with these individualized effects, baseline brain, environmental, and pubertal measures were compared between individuals with and without significant effects. Three representative behavioral pairs are shown in **Figure 4C**. Stronger structural connectivity between right posterior cingulate and right marginal cingulate regions, as well as between right postcentral and right superior precentral regions, were associated with significant effects linking lack of planning to aggressive behavior (*p* < 0.001). Higher fractional anisotropy in the right intraparietal sulcus and transverse parietal sulci was associated with significant effects linking behavioral inhibition to somatic complaints (*p* < 0.001). Individuals with moderate weekend screen time were more likely to show significant effects linking visuospatial accuracy to attention problems (*p* < 0.001).

## DISCUSSION

Cognitive and personality development is closely related to trajectories of adolescent mental health, yet these relationships arise within a broader developmental context shaped by brain maturation, environmental exposures, and pubertal transitions (17,60,61). Accordingly, the current study adopted comprehensive measures from the ABCD Study and identified multiple significant developmental pathways linking cognition and personality with mental health. Extending previous studies, we reveal substantial inter-individual variability in these associations and identify mechanisms underlying this variability depending on individual brain and environmental profiles. Among the significant behavioral associations identified through mediation and longitudinal interaction models, several behavioral traits were repeatedly involved (**Figures 2** and **3**). Personality traits, particularly behavioral inhibition and negative urgency, showed substantial involvement in these associations. Negative urgency has been associated with increased self-harm, problematic alcohol use, and disordered eating behaviors (62,63), all of which are relevant to mental disorders. Early-life behavioral inhibition, particularly in interaction with parental influences, can influence long-term mental development (64,65), including greater difficulties in mentalizing ability (66). This pattern is consistent with prior research suggesting that impulsivity-and inhibition-related traits play important roles in emotional regulation and vulnerability to psychopathology. Cognitive measures such as vocabulary were also involved in several pathways, consistent with evidence that impairment in vocabulary ability during adolescence is associated with lower engagement in education and employment, and increased risk of affective disorders (67). Taken together, these findings suggest that such behavioral traits may shape patterns of interpersonal engagement by increasing internalized stress while limiting access to externalized social support, thereby increasing vulnerability to a range of mental health problems, including depression, social difficulties, and psychosis-related symptoms (64,65,68).

At the brain level, significant effects in the identified behavioral associations were concentrated in regions within the frontal, temporal, and occipital lobes. Prior research has repeatedly implicated these regions in personality disorders and cognitive impairments, particularly in processes related to attention and impulse control (69–71). Prefrontal regions support executive functions such as planning, attentional control, and behavioral regulation (69), consistent with the involvement of traits such as lack of planning and behavioral inhibition observed in our results. Disruptions in these executive control systems may influence how individuals regulate impulsive tendencies, thereby linking personality traits with downstream mental health outcomes. Abnormalities in temporal lobe regions, particularly the superior temporal gyrus and medial temporal structures, have been associated with impairments in language-related cognitive processes as well as personality-related behavioral changes (70,72,73). These alterations may help explain the involvement of cognitive abilities such as vocabulary and personality traits observed in our analyses. In addition, altered occipital activity related to emotional visual processing has been associated with both cognitive impairment and affective dysregulation in major depressive disorder (71,74), suggesting that visual-emotional processing systems may contribute to links between behavioral traits and depressive symptoms observed in our analyses. Collectively, these findings suggest that the regions implicated in our analyses form part of neural systems supporting executive control, language and social cognition, and emotion-related behavioral regulation, which may contribute to the behavioral associations between cognitive abilities, personality traits, and mental health outcomes.

Compared to brain-derived metrics, environmental factors exhibited more widespread effects, with lifestyle-related exposures showing the strongest contribution. In particular, weekend screen use and sleep problems emerged as prominent factors. Excessive screen time has been consistently associated with poorer cognitive performance (68) and reduced well-being (75), while sleep problems, which are highly prevalent across mental disorders (76), have been linked to adverse personality traits (77) and reduced positive mood (78). Together, these lifestyle-related exposures may exert sustained influences on behavioral patterns, thereby shaping broader mental health outcomes. Family-related exposures also showed notable effects. For example, family conflict was significantly associated with poorer well-being, partly mediated by cognitive abilities such as cognitive flexibility (79), suggesting that family context may shape behavioral pathways linking cognition and mental health. In contrast, pubertal factors showed relatively limited effects on the behavioral associations. Hormonal measures were significant in only a few behavioral pairs, predominantly involving reading ability. These finding are consistent with evidence linking testosterone to brain regions involved in verbal dominance (80) and with recent findings reporting no clear causal associations between estradiol exposure and brain structure or mental health outcomes (81). Overall, these findings highlight the prominent role of environmental context in shaping behavioral associations between cognition, personality, and mental health (14,15,61).

Beyond group-wise findings, our analyses revealed substantial inter-individual heterogeneity in the associations linking cognition and personality with mental health (**Figure 4**). This pattern suggests that behavioral relationships identified at the population level do not operate uniformly across adolescents but instead depend on individual neurobiological and environmental contexts (19,25–27). For example, stronger structural connectivity between posterior and marginal cingulate regions was associated with significant effects linking lack of planning to aggressive behavior. These regions form part of large-scale networks supporting cognitive control and self-referential processing (82,83). In addition, adolescents with moderate levels of weekend screen time were more likely to show significant associations between visuospatial performance and attention problems. Screen use has been associated with alterations in attentional control and cognitive engagement, potentially through repeated exposure to rapidly changing visual stimuli and reduced opportunities for sustained attention (84,85). Moderate exposure may therefore interact with individual cognitive profiles to influence attentional functioning, whereas both very low and very high screen use may reflect different behavioral or lifestyle patterns. Together, these findings underscore the importance of considering behavioral associations from an individualized perspective.

The observed inter-individual heterogeneity in behavioral associations has potential implications for intervention strategies. Prior research has demonstrated that psychiatric symptoms can be reduced through interventions targeting cognition and personality-related processes. For example, improving attention and focus has been shown to improve social and behavioral symptoms in children with neurodevelopmental disorders (86–89). In patients with ADHD, improving attention through behavioral interventions or stimulant medication has also been linked to improved social functioning and decreased symptoms from comorbid psychopathologies (86), while in autism, joint-attentional interventions have generalized to better social communication skills (87). Likewise, our results indicate that modulating attention may represent a particularly promising target for improving thought-disorder symptoms and social functioning (**Figure 4**). Indeed, many of the cognitive, personality, and behavioral trait pairs identified here align with patterns previously reported in the literature, such as the association between stronger speech ability and reduced manic symptoms (90), or the relationship between negative urgency and poorer depression and anxiety outcomes (91). Nevertheless, not all identified pairs are currently supported by empirical evidence, necessitating further clinical investigation to validate and extend our findings. Given that neuropsychiatric treatments are typically multifactorial, developing targeted interventions focused on discrete cognitive and personality traits may provide useful complements to established approaches such as cognitive behavioral therapy, which predominantly focuses on higher-order cognitive processes (92). Overall, our findings provide data-driven guidance on the cognitive and personality traits most relevant to different mental health domains, offering a foundation for future clinical validation and the development of personalized therapies.

Several limitations should be considered when interpreting these findings. First, the analyses remain observational, and therefore the identified mediation pathways and longitudinal effects should not be interpreted as definitive causal relationships. Although longitudinal modeling strengthens temporal interpretation, unmeasured confounding factors may still influence the observed associations. Second, the longitudinal sample (*N* = 1437) was smaller than the cross-sectional sample due to attrition across follow-up assessments, which may introduce bias and reduce statistical power for change-based analyses. Third, although this study included a comprehensive set of measures available in the ABCD dataset, not all relevant behavioral or contextual factors were assessed across all time points. In particular, disruptions related to the COVID-19 pandemic limited the availability of some follow-up assessments, which may restrict the coverage of certain developmental influences. Finally, participants in the ABCD cohort were predominantly recruited from a high-income country context, which may limit the generalizability of the findings to populations in lower- and middle-income settings, where the burden of mental illness is often greater (61,93).

In conclusion, the present study provides a comprehensive framework for examining how cognitive and personality traits relate to adolescent mental health within a broader developmental context. By integrating brain structural measures, environmental exposures, and pubertal factors within longitudinal and individualized analyses, our findings reveal multiple behavioral pathways linking cognition and personality to mental health outcomes. Importantly, these associations vary substantially across individuals and are shaped by differences in neurobiological organization and environmental context. These results highlight the importance of considering individual developmental profiles when characterizing behavioral risk mechanisms and provide a foundation for future research aimed at developing more targeted and personalized strategies for mental health prevention and intervention.

## ACKNOWLEDGMENTS AND DISCLOSURES

This work was supported by the Canadian Institutes of Health Research (to SK), the CFREF/HBHL (to ACE), and the China Scholarship Council (to JY).

Data used in this study were obtained from the ABCD Study (https://abcdstudy.org), held in the NDA. The ABCD Study is a multisite, longitudinal study designed to recruit over 10,000 children aged 9-10 and follow them for 10 years. It is supported by the National Institutes of Health (NIH) and other federal partners under award numbers U01DA041048, U01DA050989, U01DA051016, U01DA041022, U01DA051018, U01DA051037, U01DA050987, U01DA041174, U01DA041106, U01DA041117, U01DA041028, U01DA041134, U01DA050988, U01DA051039, U01DA041156, U01DA041025, U01DA041120, U01DA051038, U01DA041148, U01DA041093, U01DA041089, U24DA041123, and U24DA041147. A complete list of participating sites and investigators is available at https://abcdstudy.org/consortium_members. The data analyzed in this study were obtained from ABCD Release 5.1.

The authors report no biomedical financial interests or potential conflicts of interest.

